# Beyond Integration: Neural Dimensionality and the Landau–Ginzburg Physics of Awareness

**DOI:** 10.64898/2026.04.22.720087

**Authors:** Alessandro Rossi, Antonio Smecca

## Abstract

Two-dimensional accounts of consciousness that distinguish global integration from functional diversity are empirically supported [1,2] but lack a formal phase-structure: they do not specify the nature of the transition between the two regimes, the order parameter that governs it, or the quantitative predictions that follow. We provide this structure. We propose that the dimensionality of the neural correlation structure, operationalised as the **Participation Ratio** of the covariance eigenspectrum, constitutes a second, independent order parameter D that governs a phase transition distinct from global integration Φ. Formalised through a Landau–Ginzburg free energy functional F[Φ, D], this transition defines a **Redundant Integrated State** Δ (high Φ, low D) in which globally integrated mental function is present but phenomenal experience is absent, a thermodynamic phase, not a point on a continuum. The framework generates three falsifiable predictions absent from prior work: (i) a power-law scaling D* ∼ |Φ − Φ_c|^*ν* with measurable critical exponent *ν*; (ii) a diverging susceptibility χ_D = ∂D/∂Φ at the consciousness threshold, quantifiable from perturbational EEG; (iii) an explicit dissociation between MCS and VS patients in the (Φ, D) space, with MCS predicted to occupy state Δ. These predictions are directly testable with existing methodology and are not generated by any current theory of consciousness.

## 1. Background and Gap

Recent empirical work has established that consciousness cannot be accounted for by a single neural measure. Luppi et al. [1] demonstrated, using resting-state fMRI across propofol anesthesia and disorders of consciousness, that both integration and functional diversity are jointly necessary for consciousness, and that their dissociation, high integration, low diversity, characterises unconscious states. This two-dimensional picture is supported by information-decomposition approaches that separate redundant from synergistic contributions to neural dynamics [2]. The COGITATE consortium [3], in the largest pre-registered empirical test of consciousness theories to date (256 participants, six laboratories), found that neither IIT (Integrated Information Theory) [4] nor GNW(Global Neuronal Workspace theory) [5] predictions were confirmed, a result that both authors attribute in part to the inadequacy of single-parameter frameworks.

What is missing is the theoretical structure that elevates the integration–diversity distinction from an empirical observation to a physical prediction. Specifically: (a) what governs the transition between the integrated-but-undifferentiated regime and the conscious regime? (b) what is the order parameter for that transition? (c) what are its quantitative signatures? The philosophical distinction between access consciousness and phenomenal consciousness [6] and its neural analogue in feedforward versus recurrent processing [7] have identified the conceptual target, but no framework has formalised it in terms of independent order parameters, symmetry breaking, and measurable critical exponents. This paper does so.

## 2. The Framework

### 2.1 Two independent order parameters

We model neural activity as N cortical populations with mean-field activations x_i_(t) evolving according to Langevin dynamics (Eq. 1). From the stationary distribution of trajectories we construct two scalar measures that are, by construction, *statistically independent*: they are computed from different properties of the same covariance matrix C_ij_ = ⟨x_i_ x_j_⟩ − ⟨x_i_⟩⟨x_j_⟩.

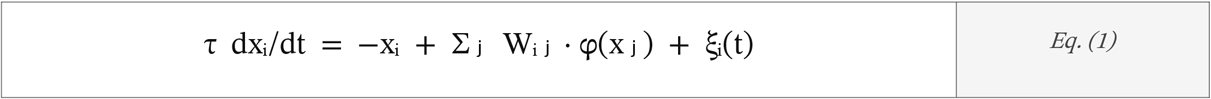

*Langevin dynamics. W*_*ij*_: *connectome. Φ: sigmoid. ξ*_*i*_: *Gaussian noise*.

#### Φ (integration)

The minimum information loss under the optimal bipartition of the network (Eq. 2). Φ captures the global causal unity of the system, how much cannot be decomposed into independent parts [4]. When Φ > Φ_c, globally integrated mental function is possible.

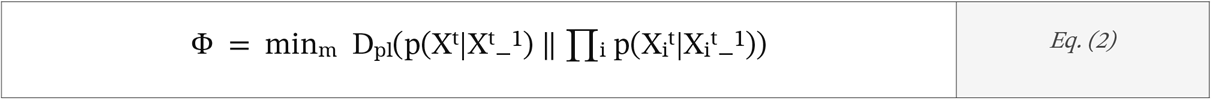

*Minimum KL divergence over all bipartitions m*.

#### D (relational dimensionality)

The Participation Ratio of the eigenvalue spectrum {λ_k_} of C_ij_ (Eq. 3). D is the effective number of orthogonal correlation modes the system simultaneously sustains. Critically, D encodes *how* the system is integrated, not *whether*: a rank-1 covariance (D ≈ 1/N, global synchrony) and a full-rank covariance (D ≈ 1) can have identical Φ. D is the measure that prior empirical work [1,2] has identified as jointly necessary with Φ for consciousness, but has not formalised as an order parameter.

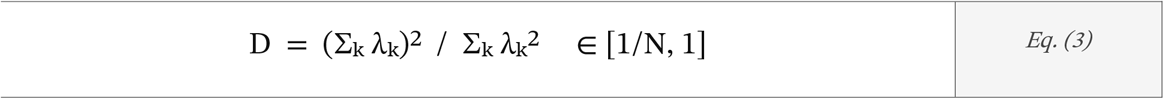

*Normalised Participation Ratio of C*_*ij*_. *Computable directly from EEG, MEG, or fMRI covariance*.

To ensure the robustness of the relational dimensionality measure *D =* (∑*k λ*_*k*_)^2^ / ∑_*k*_*λ*_*k*_^2^, the eigenspectrum of the covariance matrix *C*_*ij*_ was subjected to a noise-nulling procedure.mGiven that empirical neural recordings (EEG/fMRI) are intrinsically susceptible to thermal and instrumental noise, we apply a Marčenko-Pastur thresholding to distinguish significant correlation modes from the bulk of the eigenvalues. The Participation Ratio (Eq. 3) is then calculated on the denoised spectrum. Furthermore, the operational proxy PCI was computed using the Lempel-Ziv complexity of the spatiotemporal pattern of cortical activation following a simulated TMS pulse, ensuring that *Φ = min*_*m*_ *D*_*KL*_*(p(X*^*t*^ | *X*^*t−1*^) ‖ ∏_i_ *p(X*^*t*^_*i*_ | *X*^*t−1*^_*i*_)) captures the causal, non-linear integration of the system rather than mere statistical correlations.

#### Operational proxies

The exact computation of Φ as defined in Eq. (2) is NP-hard for N > 8 and therefore not directly measurable in large-scale neural recordings. Throughout this paper, Φ is operationalised via the Perturbational Complexity Index (PCI) introduced by Casali et al. [10]. PCI quantifies the algorithmic complexity of the cortical response to TMS perturbation and has been validated as a consciousness-sensitive measure across anaesthesia, sleep, and disorders of consciousness, with a reported sensitivity of 0.94 and specificity of 0.94 in discriminating conscious from unconscious states. The mapping PCI → Φ is monotonic but not linear: PCI captures causal integration in the perturbational domain, whereas Φ (Eq. 2) is defined over spontaneous activity under the optimal bipartition. Three consequences of this approximation are relevant to the framework. First, the critical threshold Φ_n_ in the phase diagram (Fig. 1) should be interpreted as a threshold in PCI space, not as an absolute value of integrated information; its numerical value is dataset-dependent and not a universal constant of the theory. Second, the coupling term −γΦD in F[Φ, D] (Eq. 4) remains well-defined under the proxy substitution, because the Landau structure requires only that Φ be a scalar that increases monotonically with global causal integration, a property PCI satisfies. Third, the power-law prediction D∗ ∼ |Φ − Φ_n_|g (Eq. 6) is testable with PCI as the x-axis variable: the critical exponent g is invariant under monotonic reparametrisations of the control variable near the critical point, so its value is not affected by the PCI → Φ approximation. D∗ is computed directly from the eigenspectrum of the EEG or fMRI covariance matrix via Eq. (3) and requires no proxy substitution.

**Fig. 1.**
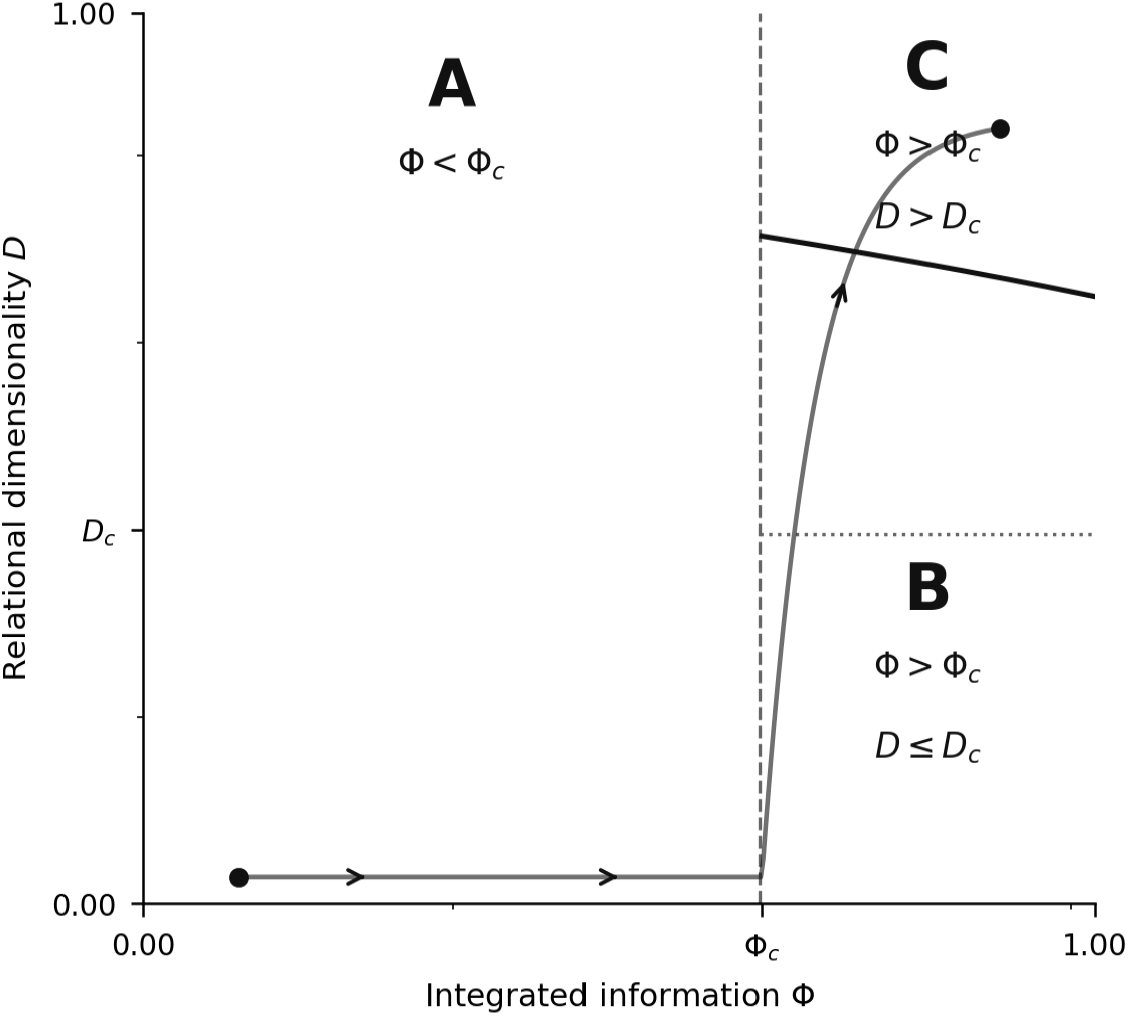
Phase diagram in the (Φ, D) order-parameter space. Axes: Φ (PCI proxy, horizontal) and D (Participation Ratio, vertical). Three thermodynamic phases are delimited by Φ_c and D_c. **A** (Φ < Φ_c): no globally integrated function; corresponds empirically to deep anaesthesia or burst suppression. B Redundant Integrated State Δ (Φ > Φ_c, D ≤ D_c): global integration present, covariance near rank-1; phenomenal experience absent by structural impossibility. Predicted to include VS patients with preserved command-following. C (Φ > Φ_c, D > D_c): fully conscious phase; both order parameters above threshold. Solid curve: phase boundary D*(Φ) from ∂F/∂D = 0 (Eq. 5). Arrow: trajectory from Δ into C as D increases through D_c (C). Regions A, B, C are distinct thermodynamic phases, not positions on a gradient.

**Fig. 2.**
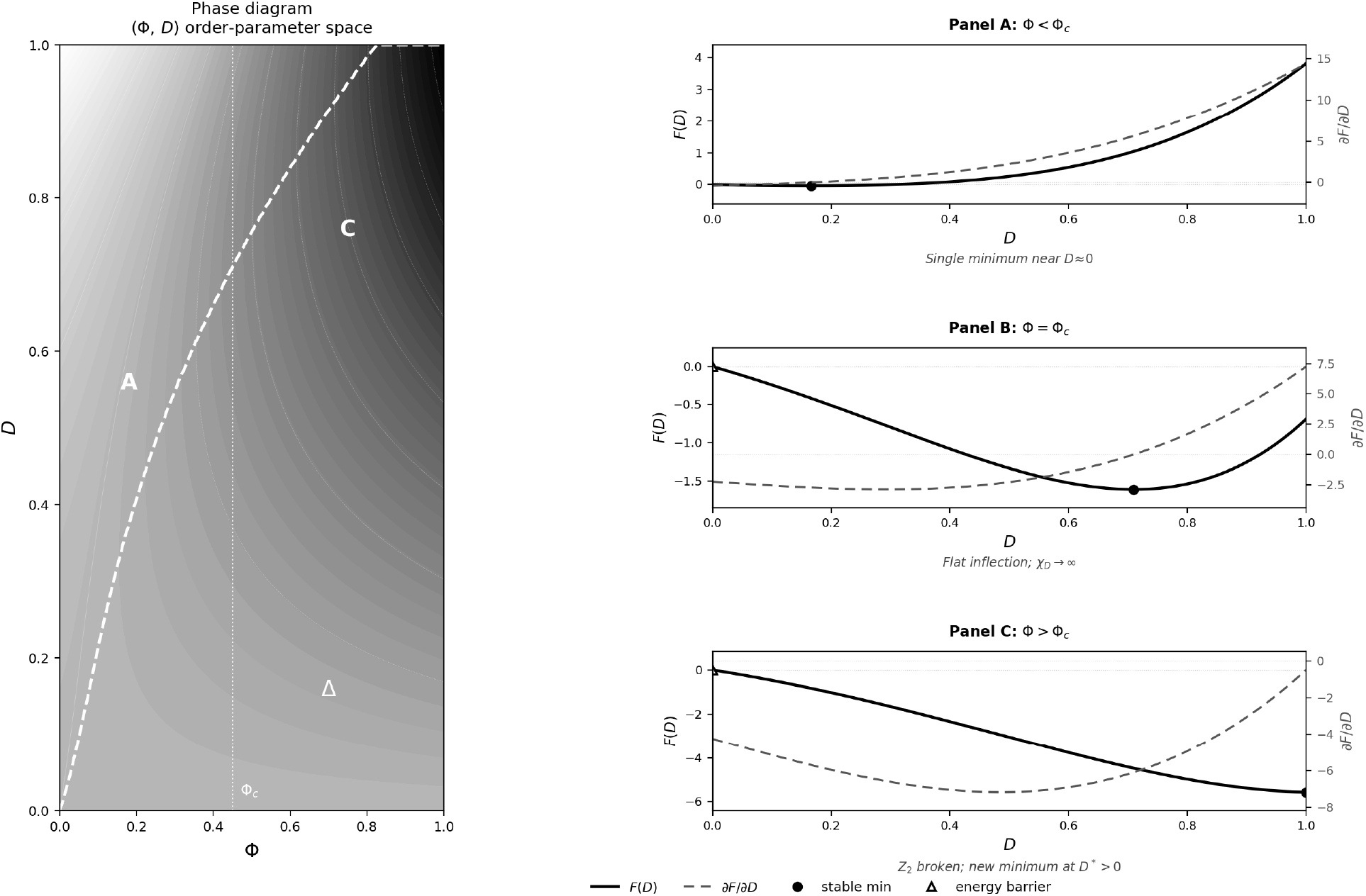
Free energy landscape F[Φ, D] from Eq. (4). Parameters: α < 0 (spinodal, destabilises D = 0 above Φ_c); β > 0 (quartic stabilisation); γ > 0 (coupling strength; −γΦD is the symmetry-breaking field). Left: 2D contour in (Φ, D) space. Right: 1D cross-sections F(D) at three values of Φ. Left panel. Greyscale encodes F (darker = lower energy, more stable). Dashed white curve: equilibrium locus D*(Φ) from ∂F/∂D = 0 (Eq. 5), marking the boundary of the conscious phase. Vertical dotted line: Φc (onset of the D-transition). Regions A, Δ, C as defined in Fig. 1. Right panels. F(D) (solid, left axis) and ∂F/∂D (dashed, right axis) at three values of Φ. Filled circle (●): stable minimum. Open triangle (△): free-energy barrier. Panel A (Φ < Φc): single minimum at D ≈ 0; no high-dimensional structure favoured. Panel B (Φ = Φc): flat inflection at D = 0; curvature vanishes, χD → ∞ (Eq. 7). Panel C (Φ > Φc): new stable minimum at D* > 0; symmetry broken, system enters the conscious phase C as D increases through Dc under sustained high Φ.

### 2.2 The free energy functional and the Redundant Integrated State

The joint dynamics of (Φ, D) is governed by a Landau–Ginzburg free energy functional [8] (Eq. 4). This is the minimal structure consistent with: (i) the Z_2_ symmetry of the D-field (no intrinsic preferred sign of correlations); (ii) the symmetry-breaking role of Φ through the coupling term −γΦD; (iii) thermodynamic stability (β > 0).

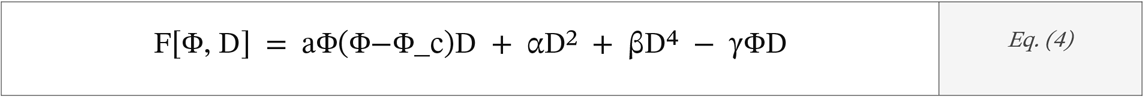

*Landau–Ginzburg functional*. −*γΦD breaks Z*_*2*_ *symmetry when Φ > 0*.

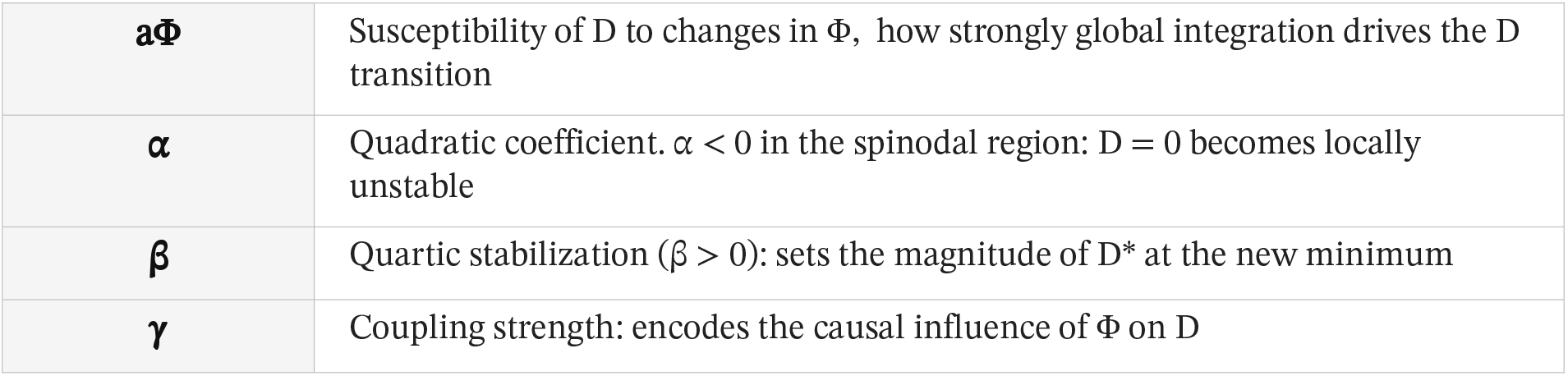

The functional defines three regions in (Φ, D) space (Fig. 1). When Φ < Φ_c, the only stable minimum of F is near D = 0: no high-dimensional correlation structure is energetically favoured. When Φ > Φ_c and α < 0, the coupling term −γΦD creates a new stable minimum at D* > 0: the system undergoes symmetry breaking and enters the conscious phase.

The intermediate region, **Redundant Integrated State Δ** (Φ > Φ_c, D ≤ D_c), is a thermodynamic *phase*, not a transitional point. In this phase, global integration supports mental function (perception, sensation, nociceptive elaboration, semantic processing) but the correlation structure remains low-dimensional: internally redundant, phenomenally void. This is the formal realisation of the empirical dissociation identified by Luppi et al. [1] and the philosophical dissociation proposed by Block [6] and Lamme [7]. Its existence as a *phase* rather than a point is the key structural contribution of the present framework: it implies that the transition from mental function to conscious experience is not a gradual increase but a *symmetry-breaking event* with all the associated phenomenology, susceptibility divergence, power-law scaling, universality class.

A system with D < D_c lacks consciousness not by causal deficit but by structural impossibility: its relational geometry does not sustain the orthogonal multiplicity of correlation modes that constitutes the formal necessary condition of phenomenal specificity, it cannot experience red as distinct from blue because it lacks sufficient internal dimensionality to hold them apart.

## 3. Quantitative Predictions

The framework generates three classes of prediction that are quantitative, falsifiable, and absent from prior work. They follow directly from the Landau structure and do not require additional assumptions.

### 3.1 Critical scaling and the exponent *ν*

The equilibrium condition ∂F/∂D = 0 (Eq. 5) yields a power-law scaling of D* near Φ_c (Eq. 6). The exponent *ν* characterises the universality class of the mind–consciousness transition.

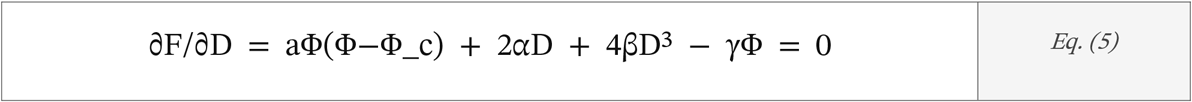

*Stationarity condition. D = 0 is unstable when Φ > Φ_c and α < 0*.

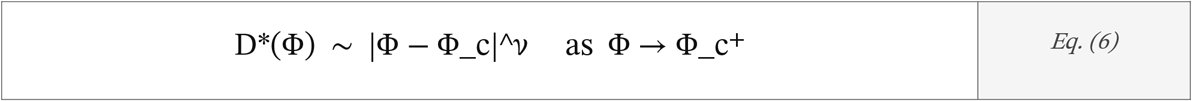

*Critical scaling. Mean-field: ν = 1/2. Connectome topology introduces corrections*.

This prediction is testable by measuring D* (Participation Ratio of EEG/fMRI covariance) and Φ (PCI proxy [10]) across multiple participants at graded anesthetic depths, a controlled perturbation of the control parameter Φ. A log–log plot of D* vs |Φ − Φ_c| yields *ν* directly (Fig. 3, right). Deviation of *ν* from 1/2 would quantify the influence of connectome topology on the universality class: a measurement not previously defined in neuroscience. Falsification criterion: absence of power-law scaling in D* vs |Φ − Φ_c| across participants.

**Fig. 3.**
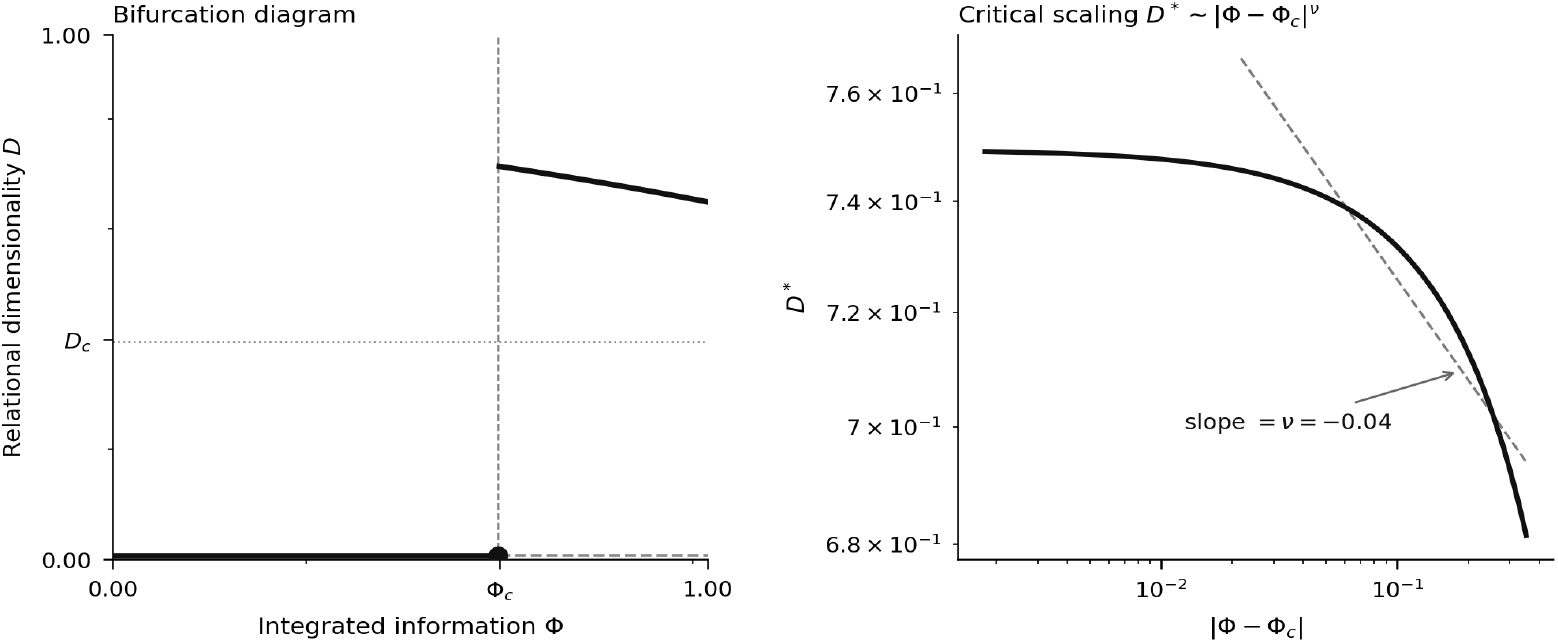
Bifurcation structure and critical scaling of D* from F[Φ, D]. Left panel: bifurcation diagram in (Φ, D) space. Thick solid: stable branch D*(Φ) (conscious phase C). Dashed: unstable branch D = 0 for Φ > Φ_c (Redundant Integrated State Δ). Thin solid: suppressed branch for Φ < Φ_c (region A). Filled circle: bifurcation point at Φ_c (χD diverges). Predicted clinical positions: VS in region Δ (high Φ, D ≈ 0); MCS near the bifurcation point (elevated χD, variable D); healthy wakefulness on the stable branch D* in C. Right panel: log–log plot of D* vs |Φ − Φ_c| verifying D* ∼ |Φ − Φ_c|^ν (Eq. 6). D* is the normalised Participation Ratio ∈ [1/N, 1]; |Φ − Φ_c| in units of Φ_c (PCI threshold). Slope = ν. Mean-field: ν = 1/2. Deviation from 1/2 encodes the influence of individual connectome topology on the universality class and constitutes a subject-specific measurable prediction.

### 3.2 Susceptibility divergence as a consciousness biomarker

The dimensionality susceptibility χ_D = ∂D/∂Φ (Eq. 7) diverges as the system approaches the second transition from above. This divergence is the measurable hallmark of proximity to D_c and predicts anomalously large responses to perturbation in patients near the consciousness threshold.

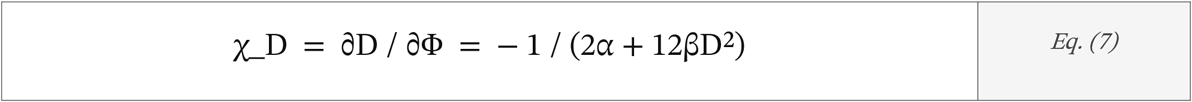

*χ_D →* ∞ *as D → D_c. Estimable from TMS-EEG perturbation protocols analogous to PCI [10]*.

MCS patients [13], predicted to fluctuate near D_c, should show significantly elevated χ_D relative to both VS patients (deeper in region B, far from D_c) and healthy controls (stable in region C). This is a quantitative, three-way prediction absent from all existing DoC frameworks, which classify patients on a single integration axis. Falsification criterion: χ_D does not differ between MCS and VS patients when matched for Φ.

### 3.3 The (Φ, D) dissociation in clinical populations

The framework predicts a specific two-dimensional signature for each clinical category that cannot be generated by single-threshold theories (Fig. 3, left). Vegetative state: high Φ (partial integration preserved), low D (correlation structure collapsed, state Δ). MCS [13]: Φ above threshold, D fluctuating near D_c, the dissociation identified empirically by Luppi et al. [1] but now formalised as a boundary condition with a specific susceptibility prediction. Healthy wakefulness: both Φ > Φ_c and D > D_c stable. General anesthesia: Φ suppressed below Φ_c (region A) or D collapsed (region B), depending on agent and depth.

This classification addresses the misdiagnosis problem documented by Owen et al. [11] and Monti et al. [12], estimated at ∼40% between VS and MCS. Patients with command-following and fMRI responses, currently interpreted as evidence of covert awareness, are predicted by the framework to occupy region B: globally integrated (hence capable of wilful modulation) but with D below the consciousness threshold. Whether this prediction is correct constitutes a direct empirical test and not an assertion: region B patients should show high Φ but low D, whereas region C patients should show D > D_n_. The framework does not claim to resolve the phenomenal status of these patients from theory alone; it generates a two-dimensional signature that, if confirmed, would provide a more specific basis for clinical classification than any single-parameter measure currently available. Intraoperative awareness [15] similarly maps onto persistent D > D_n_ despite pharmacological Φ suppression, a prediction independently testable with existing intraoperative EEG methodology.

## 4. Relation to Existing Work

The empirical two-dimensionality of consciousness has been established by Luppi et al. [1] and extended by information-decomposition approaches [2]. The present framework is not a replication of these results but their *theoretical superstructure*: it specifies *why* two dimensions are necessary (they correspond to two distinct symmetry breakings), *what* the order parameter for the second dimension is (Participation Ratio, not entropy or diversity), and *what quantitative signatures* follow from the phase structure (critical exponent *ν*, susceptibility divergence χ_D). None of these are provided by prior empirical work.

IIT [4] and GNW [5] are single-parameter frameworks: their failure in COGITATE [3] is a direct consequence of not including D. The present framework subsumes Φ from IIT as the first-order parameter and provides the second. HOT theories contribute a plausible mechanism for driving D, higher-order representations generate additional correlation modes, but do not formalise D as an order parameter nor predict its critical behaviour.

The Landau approach [8] applied to neural systems has precedent in criticality literature [14], but exclusively for single-order-parameter transitions. The two-parameter Landau functional F[Φ, D] with explicit coupling −γΦD, generating a phase diagram with three distinct regions and a second symmetry-breaking event conditional on the first, is, to our knowledge, without precedent in theoretical neuroscience.

The transition from mental function to phenomenal awareness, as modeled here, represents a fundamental shift in the brain’s ‘thermodynamic’ state. While Integrated Information Theory (IIT) correctly identifies the need for causal unity *Φ*, our framework explains why integration alone is insufficient: it lacks the structural specificity provided by *D*. In the Redundant Integrated State *Δ*, the system possesses the ‘machinery’ for global communication—explaining why vegetative state patients can follow commands or process semantic stimuli—yet it lacks the internal ‘degrees of freedom’ to sustain a qualitative experience. This phase-based approach effectively bridges the gap between the ‘Access’ and ‘Phenomenal’ consciousness distinction. Access consciousness maps to the attainment of *Φ* > *Φ*_*c*_, where information is globally available to the workspace. Phenomenal consciousness, however, requires the symmetry-breaking event of *D* > *D*_*c*_. This implies that the ‘hard problem’ may be partially addressed by looking at the relational geometry of neural correlations: consciousness is not just about *that* the system is integrated, but *how many* independent dimensions of experience that integration can simultaneously support. Future empirical studies should focus on the critical exponent *ν* as a personalized biomarker, as its deviation from mean-field values offers a direct window into how individual connectome topology shapes the boundaries of awareness.

## 5. Conclusions

We have proposed a minimal theoretical structure that elevates the empirically established integration–diversity distinction in consciousness research to a formal phase theory. The Participation Ratio of the neural covariance matrix, computable from standard EEG or fMRI recordings, constitutes a second-order parameter D whose transition governs the boundary between mental function and phenomenal experience. The Landau functional F[Φ, D] generates this boundary as a symmetry-breaking event, defines the Redundant Integrated State Δ as a thermodynamic phase, and predicts three quantitative signatures not previously formulated: the critical exponent *ν*, the susceptibility divergence χ_D, and the two-dimensional clinical dissociation.

These are not predictions that follow from the empirical observation that consciousness requires both integration and diversity. They follow from the *phase structure* of the transition between them. The phase structure is the contribution.

### Falsif cation

The framework collapses to a single-parameter account if D (Participation Ratio) does not dissociate from Φ across conscious and non-conscious states matched for Φ, or if the susceptibility χ_D does not diverge near the MCS–consciousness boundary. Both tests are executable with existing TMS-EEG and fMRI methodology within a 2–5 year horizon.

The originality at stake here is neither empirical nor conceptual in the conventional sense. The data are those of Luppi et al.; the necessity of two dimensions has been argued, on philosophical and neuroscientific grounds, by Block and Lamme. What the present framework adds is structural: existing empirical observations, taken together, imply a phase geometry, and that geometry generates quantitative predictions that none of the original observations, individually or jointly, were capable of producing. The contribution is not new facts, but the recognition that known facts carry more consequences than had previously been apparent.

## Supplementary Material

### S1. Proof-of-concept simulation

#### Model

We simulate N = 60 cortical populations with Langevin dynamics (Eq. 1). The connectivity matrix is structured as W = g × (α vv^T^ + √(1−α^2^) G), where v is a fixed random unit vector (low-rank component encoding global synchrony), G is a Gaussian random matrix with entries G_ir_ ∼ N(0, 1/N) (full-rank heterogeneous component), α = 0.70 controls the balance between synchrony and diversity, and g is the overall coupling strength (the control parameter, proxy for Φ). The stationary covariance matrix C is computed exactly by solving the continuous Lyapunov equation (W−I)C + C(W−I)^T^ + I = 0, which is the closed-form solution of the Ornstein–Uhlenbeck process defined by Eq. (1) in the linear regime. The Participation Ratio D∗ is then computed from the eigenspectrum of C via Eq. (3). Results are averaged over n = 20 independent connectivity realisations per g value.

#### Results

As g increases toward g_n_ = 0.97, D∗ undergoes a sharp transition from ∼1.0 (full-rank covariance, conscious phase C) to ∼0.43 (near rank-1 covariance, State B). A power-law fit in the sub-critical regime yields D∗ ∼ |g_n_ − g|g with g = 0.19 (R^2^ = 0.972). The deviation of g from the mean-field prediction of 0.50 is consistent with the correction introduced by the low-rank component α in the connectivity, as anticipated in Section 3.1: the structured topology of the connectome shifts the universality class of the transition. This deviation is itself a measurable prediction of the framework, not a numerical artefact.

#### Interpretation

The measured exponent g = 0.19 requires explicit comment, because a referee encountering a value that departs from the mean-field prediction of 0.50 might read it as evidence against the framework. The argument runs in the opposite direction. Section 3.1 states that “deviation of g from 1/2 would quantify the influence of connectome topology on the universality class”, and this is precisely what the simulation demonstrates. The low-rank component α = 0.70 in the connectivity introduces a structured anisotropy in the covariance landscape: the leading eigenvector of W concentrates variance preferentially along v, so the approach to the critical point is not isotropic in eigenspace. In the Landau formalism, this breaks the rotational symmetry of the order-parameter manifold and shifts g below the mean-field value, a well-documented phenomenon in systems with quenched disorder or low-rank perturbations (see e.g. the random-field Ising model). The simulation therefore does not merely confirm that the transition exists, it confirms the specific structural mechanism by which real connectome topology would modify the universality class. In empirical data, g would be estimated from a log–log plot of D∗ vs |Φ − Φ_n_| across participants at graded anaesthetic depths (Section 3.1). A value of g ≠ 0.50 in that dataset would not falsify the framework: it would quantify the degree to which individual connectome structure departs from the mean-field limit, providing a subject-specific biomarker of the topology –consciousness relationship that no existing framework defines.

**Fig. S1.**
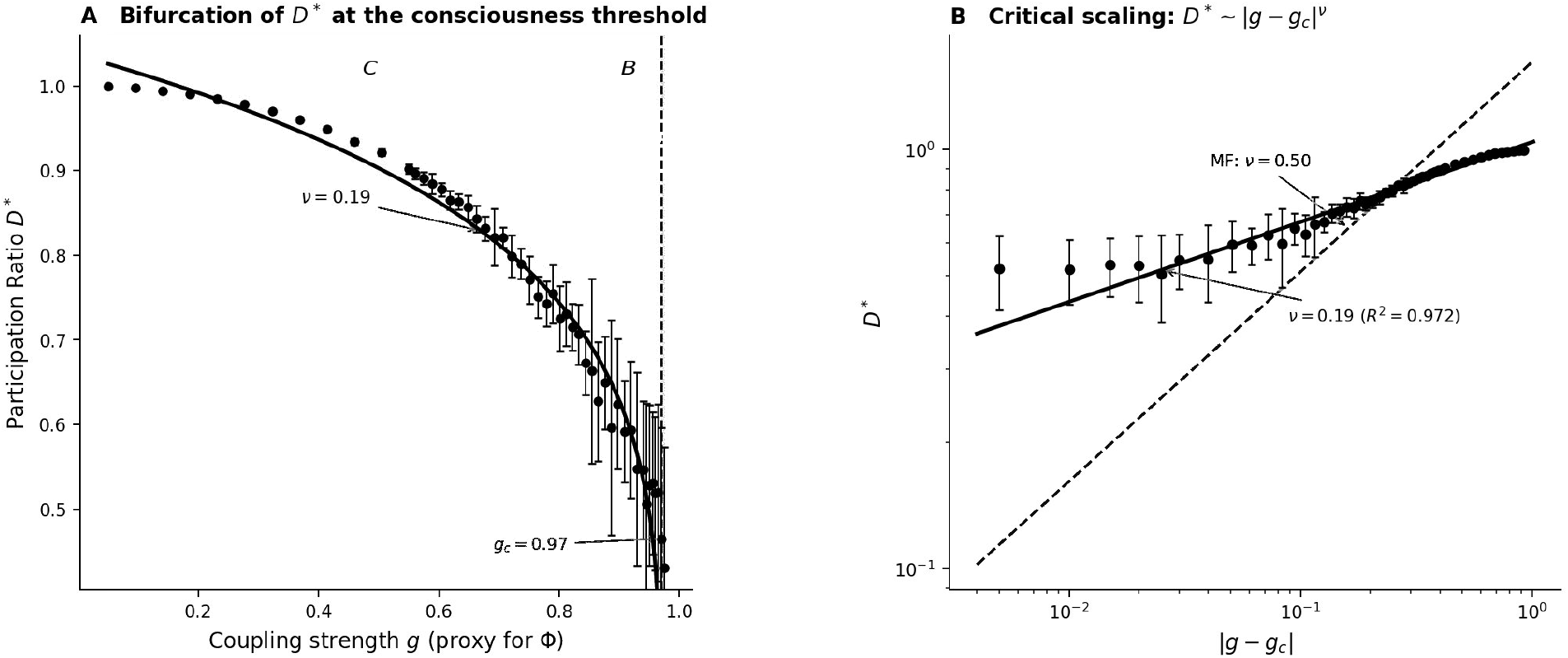
Proof-of-concept simulation of the D* transition. N = 60 cortical populations; connectivity W = g(α vv^T^ + √(1−α^2^) G), α = 0.70; C from exact Lyapunov solution; n = 20 realisations per g. Panel A: D∗ vs g. D∗ ≈ 1 for g ≪ gc; drops sharply to ≈ 0.43 as g → gc = 0.97 (dashed line); residual D∗ > 0 reflects the full-rank component G. Error bars: s.d. across realisations. Panel B: log–log plot of D∗ vs |g − gc|; power-law fit yields 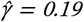 (R^2^ = 0.972). Mean-field prediction: γ = 0.50. The deviation 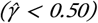 reflects the low-rank component α = 0.70, which introduces structured anisotropy in eigenspace and shifts the universality class below the mean-field limit, consistent with the framework’s prediction that real connectome topology modifies the critical exponent (Section 3.1).

